# Antifungal activity and mechanisms of D-limonene against *Fusarium oxysporum*, a pathogen of potato dry rot

**DOI:** 10.1101/2025.10.06.675995

**Authors:** Qianhao Xia, Pan Dong

## Abstract

**BACKGROUND:** Potato dry rot (PDR), caused by *Fusarium* species, seriously threatens potato production and postharvest quality. Although limonene has demonstrated antifungal activity, previous studies have mainly focused on limonene-containing essential oils or phenotypic growth inhibition, while its cellular and molecular mechanisms and application potential against PDR remain insufficiently understood.

**RESULTS:** D-limonene inhibited the growth of *Fusarium oxysporum* in a concentration-dependent manner, with a half-maximal inhibitory concentration (IC₅₀) of 8.32 μL/mL. It also altered hyphal morphology, reduced pathogenicity and spore germination, decreased spore viability, and changed the chitin-associated fluorescent brightener 28 staining pattern. Transcriptome analysis identified 1,884 differentially expressed genes, including 1,027 downregulated and 857 upregulated genes. Kyoto Encyclopedia of Genes and Genomes analysis and targeted annotation screening revealed extensive remodeling of cell-wall-related processes. Gene Set Enrichment Analysis further indicated suppression of ergosterol biosynthesis and ribosome biogenesis. D-limonene also increased sensitivity to temperature, salinity, and oxidative stresses and showed additive and synergistic interactions with mancozeb and hymexazol, respectively.

**CONCLUSION:** Our results reveal that D-limonene shows potential as a bio-based component for the integrated management of PDR, providing a theoretical basis for its application.

## 1. Introduction

Potato dry rot (PDR) is a widespread postharvest disease caused by several *Fusarium* species and represents a major threat to global potato production and postharvest quality.^1^ Among the causal agents, *Fusarium oxysporum* is one of the prevalent and virulent pathogens. It was first isolated from potato vascular tissue by Weiss in 1924 and can cause vascular wilt during plant growth as well as dry rot in stored tubers.^2–4^ Several strategies have been developed to manage PDR, including the use of disease-free seed tubers, early pathogen detection, improved storage conditions, and biological control.^5–7^ Nevertheless, chemical fungicides such as 2-aminobutane, imazalil, flusilazole, and difenoconazole remain among the most effective control measures.^8^ Their extensive and long-term use, however, may promote fungicide resistance, leave residues on agricultural products, and cause environmental contamination.^5,9,10^ Therefore, environmentally friendly alternatives are urgently needed for PDR management. Limonene (C10H16) is a cyclic monoterpene containing a six-membered carbon ring and two carbon–carbon double bonds. It occurs as two enantiomers, (R)-limonene (D-limonene) and (S)-limonene.^11^ As a plant-derived hydrocarbon, limonene undergoes aerobic microbial biodegradation under environmental conditions.^12^ It has been widely used in foods and cosmetics ^13^ and as a renewable solvent and chemical feedstock.^11,13^ In crop protection, limonene has demonstrated insecticidal and repellent activities against insects,^14–18^ fumigant toxicity against stored-product insects,^22^ and acaricidal activity against mites.^19–21^ Its potential as a botanical pesticide has also been explored.^23^ Limonene has shown antimicrobial activity against postharvest pathogens such as *Colletotrichum* ^25^ and several yeast species, including *Candida albicans*, *Zygosaccharomyces rouxii*, *Candida parapsilosis*, and *Candida tropicalis*.^26–29^

Among phytopathogenic fungi, evidence from *Fusarium* species particularly supports the antifungal and antimycotoxigenic potential of limonene. It inhibited *F. proliferatum* by damaging cell membranes and by reducing H3K9ac and H3K27ac, thereby suppressing ribosomal function, mitochondrial biosynthesis, and other metabolic processes.^30^ Limonene also reduced fumonisin B1 biosynthesis in *F. verticillioides*,^31^ while a limonene formulation suppressed both mycelial growth and deoxynivalenol production in *F. graminearum*.^32^ These studies collectively indicate that limonene can interfere with *Fusarium* growth, membrane integrity, cellular metabolism, ribosome-related processes, and mycotoxin biosynthesis.^30–32^ Although purified (S)-limonene has been shown to inhibit the mycelial growth and extracellular enzyme activities of *F. oxysporum*,^33^ direct evidence concerning the antifungal activity and mechanisms of D-limonene against this pathogen remains limited. Given that D-limonene is the predominant naturally occurring enantiomer in citrus peel oils and is more widely available for commercial and agricultural applications,^11^ systematic investigation of its antifungal activity against *F. oxysporum* is particularly warranted. Therefore, this study combined phenotypic, pathogenicity, cytological, transcriptomic, stress-sensitivity, and fungicide-combination analyses to characterize the antifungal activity and molecular responses of a PDR-associated *F. oxysporum* strain to D-limonene and to assess its potential for the integrated management of PDR.

## 2. Materials and Methods

### 2.1 Materials

The *F. oxysporum* strain used in this study was the same strain described by Ren et al. and was cultured on potato dextrose agar (PDA) medium or in potato dextrose broth (PDB) at 28 °C in the dark according to their reported method. ^34^ D-Limonene [(R)-limonene; purity 99%; CAS No. 5989-27-5] and DMSO were purchased from Shanghai Macklin Biochemical Co., Ltd. (Shanghai, China). Hymexazol and mancozeb were obtained from Shandong Kunniu Plant Protection Co., Ltd. (Shandong, China) and Shandong Lebang Chemical Co., Ltd. (Shandong, China), respectively. Healthy, uniform-sized, and firm potato tubers were obtained from a local market for subsequent experiments.

### 2.2 Effect of D-Limonene on *F. oxysporum* Hyphal Growth

PDA medium was prepared as described in Section 2.1. D-Limonene was emulsified in 0.3 mL DMSO and added to PDA medium at 60 °C to obtain final concentrations of 40.0, 20.0, 10.0, 5.0, 2.5, and 0 μL/mL, with a final volume of 30 mL. The medium was then poured into petri plates at 10 mL per plate. A 6-mm-diameter mycelial plug taken from the edge of a 5-day-old *F. oxysporum* colony was placed at the center of each plate. The 0 μL/mL D-limonene + DMSO group was used as CK. Each group included three biological replicates. Plates were incubated at 28 °C for 5 d, after which colony diameters were measured using the cross-measurement method. The inhibition rate (%) was calculated as follows: inhibition rate (%) = [(C − T) / (C − D)] × 100, where C, T, and D represent the control colony diameter, treated colony diameter, and disc diameter, respectively.^34^ GraphPad Prism 10 software was used to plot the inhibition curve and calculate the IC₅₀ value.

### 2.3 Effect of D-Limonene on *F. oxysporum* Hyphal Morphology

PDB medium was prepared as described in Section 2.1. D-Limonene was emulsified in 0.2 mL DMSO and added to PDB medium at 60 °C to obtain the IC₅₀ concentration, with a final volume of 20 mL. CK consisted of 20 mL PDB containing 0.2 mL DMSO only. Spores were harvested from 5-day-old *F. oxysporum* cultures on PDA plates with sterile water, counted using a hemocytometer, diluted to 10⁶ spores/mL. An aliquot of the resulting spore suspension was then inoculated into PDB medium. Each group included three biological replicates. After 5 d of incubation at 28 °C, mycelia were collected, mounted on slides, and observed under a microscope (DM IL LED, Leica Microsystems, Germany). For cold field-emission scanning electron microscopy (cold FE-SEM), A 6-mm-diameter mycelial plug from the edge of a 5-day-old *F. oxysporum* colony was placed at the center of each PDA medium plates supplemented with or without D-limonene at the IC₅₀ concentration, and then incubated at 28 °C for 3 d. Subsequently, mycelia were collected and observed using a SU8600 cold FE-SEM (Hitachi, Tokyo, Japan).

### 2.4 Effect of D-Limonene on the Pathogenicity of *F. oxysporum*

*F. oxysporum* was cultured on PDA plates supplemented with D-limonene at 0, 2.5, 5, 10, 20, and 40 μL/mL for 5 d. A 6-mm-diameter mycelial plug from each culture was placed at the center of each potato slice (4 cm ×3 cm ×0.7 cm) and incubated at 25 °C under humid conditions for 5 d. Lesion diameters were measured using the cross-measurement method, and inhibition curves were plotted using GraphPad Prism 10 software.

### 2.5 Effect of D-Limonene on the Germination Rate of *F. oxysporum*

Spore suspension prepared as described in Section 2.3 was evenly spread onto the PDA medium supplemented with or without D-limonene at the IC₅₀ concentration and incubated at 28 °C in the dark for 6 h. Spore germination was observed under the same microscope in Section 2.3 and was defined as a germ tube length reaching half the spore length. Germinated and total spores were counted using the five-point sampling method. The germination rate was calculated as follows: germination rate (%) = (number of germinated spores / total number of spores) ×100. Each treatment included three biological replicates.

### 2.6 Effect of D-Limonene on the Spore Viability of *F. oxysporum*

Spore suspension prepared as described in Section 2.3 was inoculated into the PDB medium supplemented with or without D-limonene at the IC₅₀ concentration, and then incubated on a rotary shaker at 28 °C for 10 h. The cultures were centrifuged at 6000 rpm for 5 min, and the pellets were resuspended in sterile water. Fluorescein diacetate (FDA) (10 μg/mL) and propidium iodide (PI) (5 μg/mL) were used to stain the spores in the dark for 15–20 min and 5 min, respectively. FDA-positive and PI-positive spores were observed under the same microscope in Section 2.3. Staining rates were calculated using the five-point sampling method as follows: staining rate (%) = (number of stained spores / total number of spores) ×100. Each treatment included three biological replicates.

### 2.7 Effect of D-Limonene on Chitin Distribution in the Cell Walls of *F. oxysporum*

Spore suspensions prepared as described in Section 2.3 were inoculated to PDB medium, and then incubated at 28 °C in the dark. When the spore germination rate reached 85%, D-limonene at the IC₅₀ concentration was added, with DMSO serving as the solvent control. After further incubation at 28 °C for 24 h, mycelia were stained with fluorescent brightener 28 (FB 28) at a final concentration of 1 mg/mL for 5 min and observed under the microscope described in Section 2.6. Each treatment included three biological replicates.

### 2.8 Effect of D-Limonene on the Stress Sensitivity of *F. oxysporum*

*F. oxysporum* was cultured on PDA medium under five stress conditions: high temperature (37 °C for 24 h), low temperature (4 °C for 24 h), ultraviolet (UV) irradiation for 30 min, high salinity (0.3 M NaCl), and oxidative stress (0.5 mM H₂O₂). Four treatments were established for each stress condition: CK, D-limonene at the IC₅₀ concentration, stress alone, and stress combined with D-limonene. After stress treatment, plates were incubated at 28 °C in the dark for 5 d. Colony diameters were measured using the cross-measurement method, and inhibition rates were calculated as described in Section 2.2. GraphPad Prism 10 software was used for plotting. Each treatment included three biological replicates.

### 2.9 Synergistic Effects of D-Limonene and Chemical Fungicides

Hymexazol and mancozeb were selected to evaluate their combined antifungal effects with D-limonene. The IC₅₀ values of hymexazol and mancozeb against *F. oxysporum* were calculated On the basis of the same method described in Section 2.2. For combination assays, CK, D-limonene, fungicide, and D-limonene + fungicide treatments were prepared, with both D-limonene and fungicide applied at their IC₅₀ concentrations. After incubation at 28 °C for 5 d, colony diameters were measured, and inhibition rates were calculated as described in Section 2.2. Agent interactions were evaluated using the Jin formula:^38^ Q = E(A+B) / (EA + EB – EA ×EB), where E(A+B), EA, and EB represent the inhibition rates of the combined treatment, agent A alone, and agent B alone, respectively. Q < 0.85, 0.85 ≤ Q < 1.15, and Q ≥ 1.15 indicate antagonistic, additive, and synergistic effects, respectively. Each treatment included three biological replicates.

### 2.10 Transcriptome Sequencing Analysis of *F. oxysporum* Following D-Limonene Treatment

*F. oxysporum* spores were inoculated into PDB supplemented with D-limonene at the IC₅₀ concentration or an equal volume of DMSO as the control. Cultures were incubated in a con-stant-temperature shaking incubator (BSD-TX270, Boxun, Shanghai, China) for 5 d. Nine culture tubes were prepared for each treatment and pooled into three independent biological replicates prior to RNA sequencing. Mycelia were harvested by centrifugation, washed with sterile water, and collected for RNA extraction. Transcriptome sequencing was performed by Tsingke Biotechnology Co., Ltd. (Chongqing, China). Clean reads were aligned to the *F. oxysporum* reference genome (http://fungi.ensembl.org/F._oxysporum/Info/Index), and transcript abundance was quantified using StringTie v2.2.1. Differential expression analysis was conducted using the DESeq2 package in R v4.4.1. Genes with an adjusted *P* value < 0.05 and |log₂ fold change (log₂FC)| ≥ 1 were considered differentially expressed genes (DEGs). Functional enrichment analyses, including Gene Ontology (GO), Kyoto Encyclopedia of Genes and Genomes (KEGG), and Gene Set Enrichment Analysis (GSEA), were performed using the clusterProfiler package in R v4.4.1. Protein–protein interaction networks (PPINs) were constructed by identifying homologous proteins in the STRING database through BLAST searches and subsequently analyzing their interaction relationships.

### 2.11 Reverse transcription quantitative PCR (RT-qPCR)

The reliability of the RNA-seq data was validated by RT-qPCR. Mycelia of *F. oxysporum* were collected using the same procedure described in Section 2.10. Total RNA was extracted using the Omega Fungal RNA Kit (R6840, Omega Bio-tek) and reverse-transcribed using the PrimeScript™ RT Reagent Kit (Takara, Japan). RT-qPCR was performed using TB Green® Premix Ex Taq™ II (RR820A, Takara) on a Bio-Rad CFX96 Real-Time PCR System. Ten DEGs (five upregulated and five downregulated) were randomly selected for RT-qPCR validation. Gene-specific primers were designed using Primer 6 and are listed in Table S4.

GAPDH was used as the internal reference gene. All experiments were conducted with three biological replicates. Relative gene expression levels were calculated using the 2^-ΔΔCt method, and log₂FC were subsequently determined for comparison with the RNA-seq results.

## 3. Results

### 3.1 D-Limonene Inhibits *F. oxysporum* Growth

After 5 d of incubation, D-limonene inhibited the mycelial growth of *F. oxysporum* in a con-centration-dependent manner (Figure 1A,C,D). The mean colony diameter decreased from 5.51 cm in the CK group to 3.23 cm at 10 μL/mL and 2.13 cm at 40 μL/mL D-limonene, corresponding to inhibition rates of 60.61% and 87.60%, respectively. Nonlinear regression analysis of the concentration–response curve using GraphPad Prism 10 yielded an IC₅₀ value of 8.32 μL/mL.

**Figure 1.**
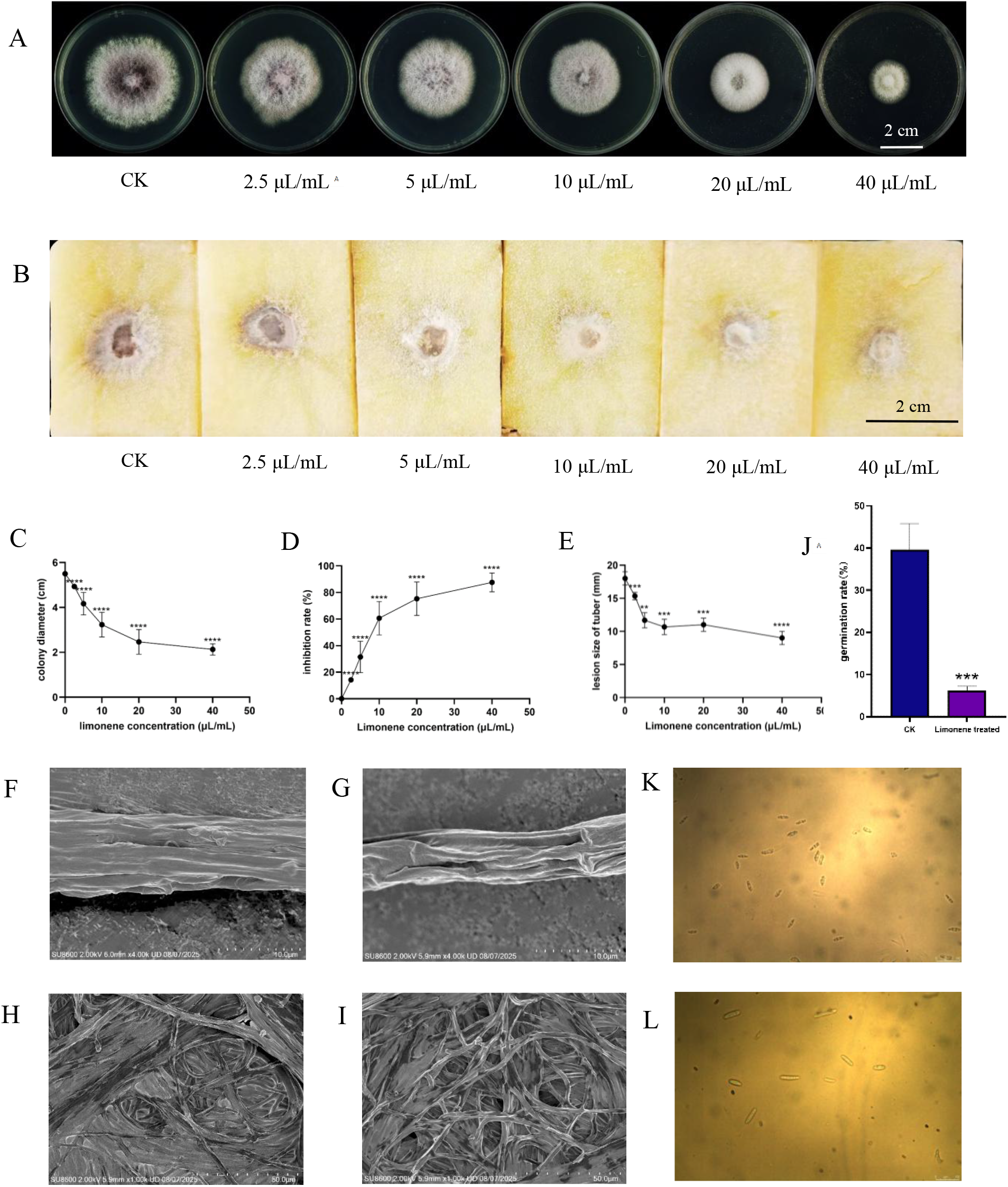
Phenotypic analysis of *F. oxysporum* under D-limonene treatment. (A) *F. oxysporum* growth after 5 d of incubation under different D-limonene concentrations. (B) Incidence of *F. oxysporum* infection on potato slices treated with different D-limonene concentrations after 5 d. (C) Colony diameter of *F. oxysporum* cultured on media containing different D-limonene concentrations for 5 d. (D) Inhibition rate of mycelial growth after 5 d under different D-limonene concentrations on *F. oxysporum*. (E) Lesion area caused by *F. oxysporum* on potato slices under different D-limonene concentrations after 5 d. (F,G) Mycelia treated with CK and half-maximal inhibitory concentration (IC₅₀) of D-limonene, observed by 10 μm cold field-emission scanning electron microscopy (cold FE-SEM). (H,I) Surface of fungal membrane treated with CK and D-limonene (IC₅₀), photographed by 50 μm cold FE-SEM. (J) Inhibition rate of spore germination of CK and the treatment group after 6 h. Each experiment was repeated at least three times, and the data are the average ±standard deviation. The asterisk (*) indicates significant differences according to the t-test (\*\**P < 0.01*; \*\*\**P < 0.001*; \*\*\*\**P < 0.0001*). (K,L) Spore germination in the D-limonene-treated (IC₅₀-treated) and untreated group after 6 h of cultivation, observed under a 40×magnification. In all figure panels, “limonene” refers specifically to D-limonene. All DMSO solvent concentrations used in this study were pre-verified to have no significant effect on *F. oxysporum* growth (Figure S1).

### 3.2 Effect of D-Limonene on the Pathogenicity of *F. oxysporum*

After 5 d, the mean lesion diameter in the CK group was 18.17 mm (Figure 1B). With increasing D-limonene concentrations from 2.5 to 5 and 10 μL/mL, the lesion diameter decreased in a concentration-dependent manner, reaching 10.67 mm at 10 μL/mL. However, further increasing the concentration to 20 and 40 μL/mL did not cause any substantial additional reduction, with the lesion diameter remaining at approximately 11 mm (Figure 1E). It should be noted that since the inoculated agar plug had a diameter of 6 mm, the measured diameter included the plug itself, indicating an even smaller actual expansion area. These results suggest that D-limonene effectively suppresses the pathogenicity of *F. oxysporum* at relatively low concentrations (≤10 μL/mL), with the inhibitory effect reaching a plateau at higher concentrations.

### 3.3 D-Limonene Affects *F. oxysporum* Hyphal Morphology

Macroscopic observation showed that D-limonene treatment altered both the texture and pigmentation of *F. oxysporum* colonies. Compared with the pale pink, relatively loose colonies in the CK group, D-limonene-treated colonies appeared whitish, denser, and thicker, with the change becoming more apparent at higher concentrations (Figure 1A). Cold FE-SEM further showed that CK hyphae were turgid, smooth, and regularly arranged, whereas hyphae exposed to D-limonene at 8.32 μL/mL (IC₅₀) were thinner and shriveled, with wrinkled surfaces, twisting, and entanglement (Figure 1F–I).

### 3.4 D-Limonene Reduces the Spore Germination Rate of *F. oxysporum*

After 6 h post-inoculation, the spore germination in the CK group reached 39.63%, whereas treatment with D-limonene at 8.32 μL/mL (IC₅₀) dramatically reduced germination to only 6.17%, representing an inhibition rate of 84.4% (*P < 0.0001*; Figure 1J–L). This result clearly demonstrates that D-limonene exerts a potent and significant inhibitory effect on *F. oxysporum* spore germination at the tested concentration.

### 3.5 D-Limonene Reduces Spore Viability of *F. oxysporum*

PI stains dead cells with compromised membrane integrity, whereas FDA stains viable cells. Cell viability was assessed by counting PI-positive (dead) and FDA-positive (live) spores in the control and D-limonene treatment groups. In the control group, the majority of spores were FDA-positive (82%), whereas only a minor proportion (less than 10%) PI-positive (Figure 2A–C,I,J). In contrast, D-limonene-treated spores were predominantly PI-positive (60%), whereas few FDA-positive spores (36%) were observed (Figure 2D–F,I,J). These results indicate that D-limonene significantly (*P* < 0.0001) reduced spore viability.

**Figure 2.**
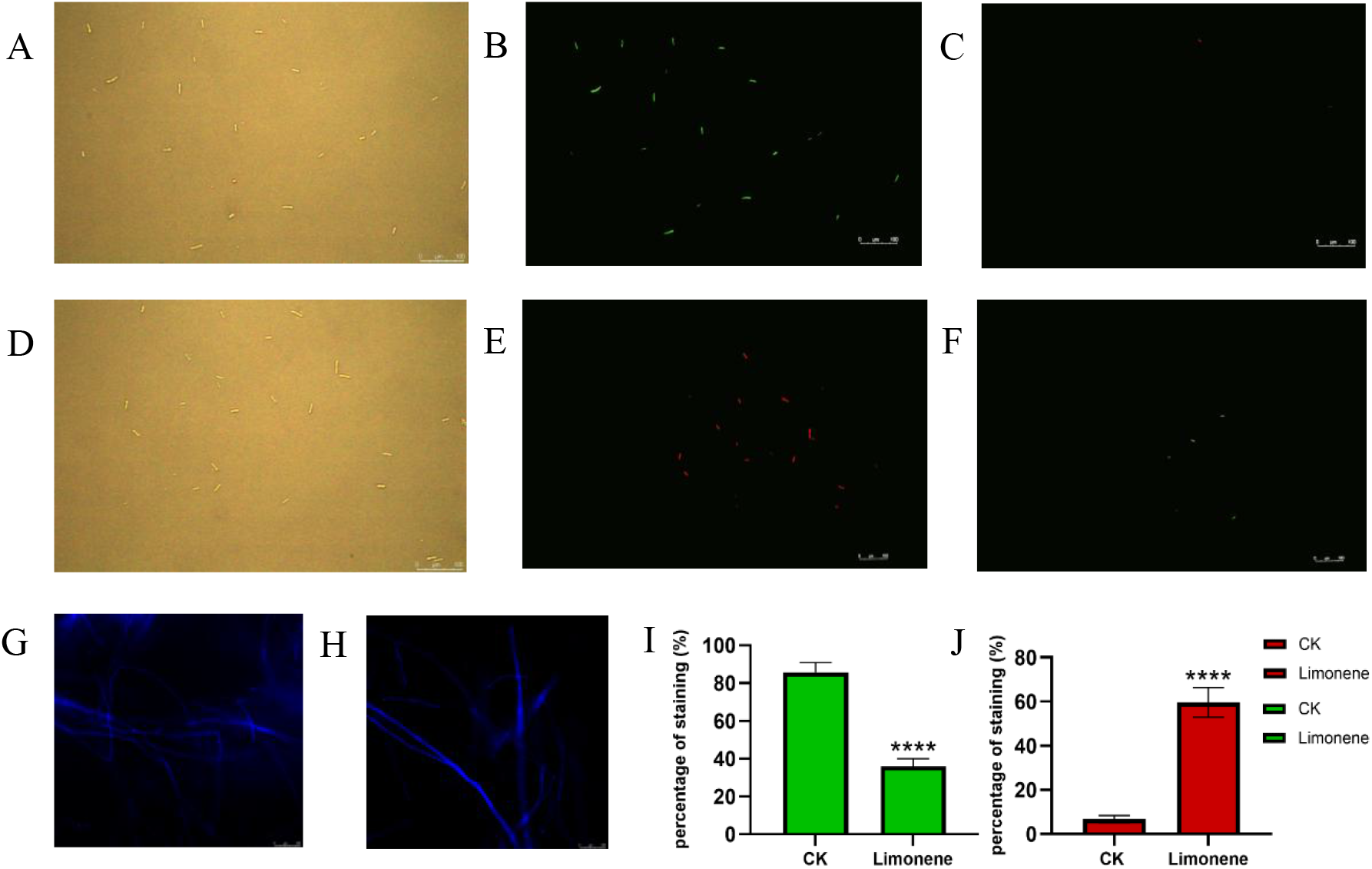
Cytological analysis of *F. oxysporum* spores and mycelia. (A,D) Bright-field images showing the total conidia in the control and D-limonene-treated (IC₅₀) groups, respectively. (B,F) FDA-stained spores in the CK and D-limonene-treated (IC₅₀) groups, respectively. (C,E) PI-stained spores in the CK and D-limonene-treated (IC₅₀) groups, respectively. (G,H) Mycelium fluorescence staining by FB28 in the CK and D-limonene-treated groups, respectively. (I,J) Percentage of FDA-positive and PI-positive spores, respectively. All experiments were repeated independently at least three times, and data are presented as mean ±standard deviation. The asterisk (*) indicates a significant difference according to a t-test (\*\*\*\**P < 0.0001*). In all figure panels, “limonene” refers specifically to D-limonene.

### 3.6 D-Limonene Alters the FB 28 Staining Pattern in the Cell Walls of *F. oxysporum*

FB 28, which specifically binds to chitin in fungal cell walls, was used to visualize chitin distribution in *F. oxysporum* hyphae. In the CK group, blue fluorescence was predominantly localized at the septa. By contrast, D-limonene-treated hyphae exhibited a distinctly altered staining pattern, characterized by significantly stronger and more diffuse fluorescence along the entire hyphal walls, rather than being restricted to the septa (Figure 2G,H). These results indicate that D-limonene treatment disrupts the normal chitin distribution of *F. oxysporum*, leading to aberrant chitin accumulation along the cell walls.

### 3.7 Effect of D-Limonene on the Partial Stress Resistance of *F. oxysporum*

Five abiotic stresses were evaluated, including low temperature, high temperature, ultraviolet (UV) irradiation, high salinity, and oxidative stress (Figure 3A–C). D-Limonene treatment (IC₅₀ = 8.32 μL/mL) alone produced an inhibition rate of approximately 29%. Among the individual stresses, high temperature (37 °C) showed the strongest inhibitory effect, with an inhibition rate of approximately 39%, whereas UV irradiation showed the weakest effect, at approximately 4%. Combining D-limonene with low temperature, high temperature, high salinity, or H₂O₂ increased the inhibition rates to approximately 38%, 63%, 40%, and 35%, respectively. The D-limonene plus high-temperature treatment produced the highest inhibition rate among all treatments. In contrast, the inhibition rate of D-limonene combined with UV irradiation was approximately 28%, similar to that of D-limonene alone. These results indicate that D-limonene increased the sensitivity of *F. oxysporum* to temperature, salinity, and oxidative stresses, with the strongest effect observed under high-temperature stress.

**Figure 3.**
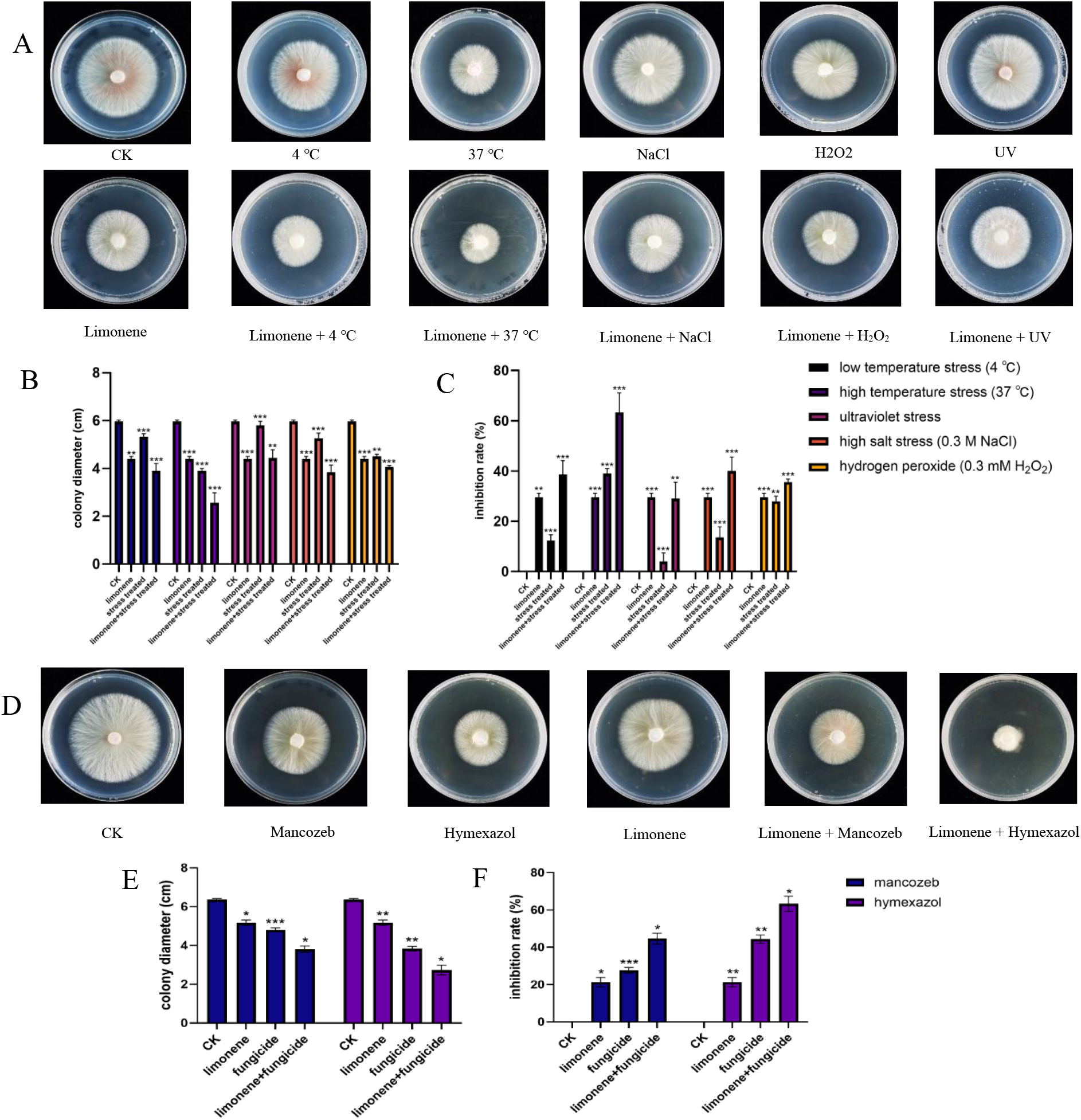
Analysis of application potential through combining D-limonene with stresses and chemical fungicides. (A–C) Colony morphology, diameter and growth inhibition rate of *F. oxysporum* under different stress conditions (low temperature, high temperature, ultraviolet (UV) irradiation, high salinity, and oxidative stress). (D–F) Colony morphology, diameter, and growth inhibition rate of *F. oxysporum* under different fungicide treatments (hymexazol and mancozeb were applied at 53.60 and 102.78 μg/mL). Each process was repeated independently at least three times, and data are the average ±standard deviation. The asterisk (*) indicates a significant difference according to a t-test (\**P* < 0.05; \*\**P* < 0.01; \*\*\**P* < 0.001). In all figure panels, “limo-nene” refers specifically to D-limonene.

### 3.8 Additive or Mildly Synergistic Effects of D-Limonene Combined with Chemical Fungicides

Two commonly used fungicides, hymexazol and mancozeb, were selected to evaluate their combined antifungal activity with D-limonene. The IC₅₀ values of hymexazol and mancozeb against *F. oxysporum* were determined to be 53.60 and 102.78 μg/mL, respectively, consistent with previously reported values.^35–37^ In PDA medium, D-limonene–fungicide combinations inhibited *F. oxysporum* more strongly than agents alone (Figure 3D,E). The inhibition rate increased from 22% with D-limonene alone and 27% with mancozeb alone to 46% with their combination; similarly, hymexazol-induced inhibition increased from 44% to 65% when combined with D-limonene (Figure 3F). Jin formula analysis ^38^ yielded Q-values of 1.068 (0.85 ≤ Q < 1.15) for D-limonene–mancozeb and 1.154 (slightly above the threshold of 1.15) for D-limonene–hymexazol, indicating additive and mildly synergistic effects, respectively. These results suggest that D-limonene can enhance the antifungal efficacy of selected fungicides, which may offer a promising strategy for optimizing fungicide application and potentially reducing dosages in agricultural practice.

### 3.9 Analysis of *F. oxysporum* Transcriptome Sequencing Results Following D-Limonene Treatment

#### 3.9.1 Global Transcriptomic Changes

Compared to the DMSO control group, a total of 1884 significantly differentially expressed genes (DEGs) were identified in D-limonene-treated *F. oxysporum*, comprising 1027 downregulated and 857 upregulated genes (*P <* 0.05, Figure 4A). Box plot analysis revealed an overall reduction in global gene expression in the D-limonene-treated samples relative to the control (Figure 4B). To validate the reliability of the RNA-seq data, 5 upregulated genes and 5 downregulated genes were randomly selected for RT-qPCR analysis. The expression trends of 9 out of the 10 selected genes were consistent with the RNA-seq results (R^2^ = 0.73, Figure 4H), supporting the reliability of the transcriptome dataset. Gene Ontology (GO) enrichment analysis identified the top five significantly enriched terms: RNA polymerase II DNA-binding transcription factor activity, transmembrane transport, RNA polymerase II cis-regulatory sequence-specific DNA binding, DNA-templated transcription, and specific sequence DNA binding (*P <* 0.05, Figure 4C). Meanwhile, Kyoto Encyclopedia of Genes and Genomes (KEGG) pathway analysis showed that 17 of 20 significantly enriched pathways were associated with metabolism, primarily involving carbohydrate metabolism and amino acid metabolism pathways (*P <* 0.05, Figure 4D). Protein–protein interaction network (PPIN) analysis further revealed a general downregulation trend across most interaction networks, with ribosome assembly-related interactions showing pronounced suppression (Figure 4E).

**Figure 4.**
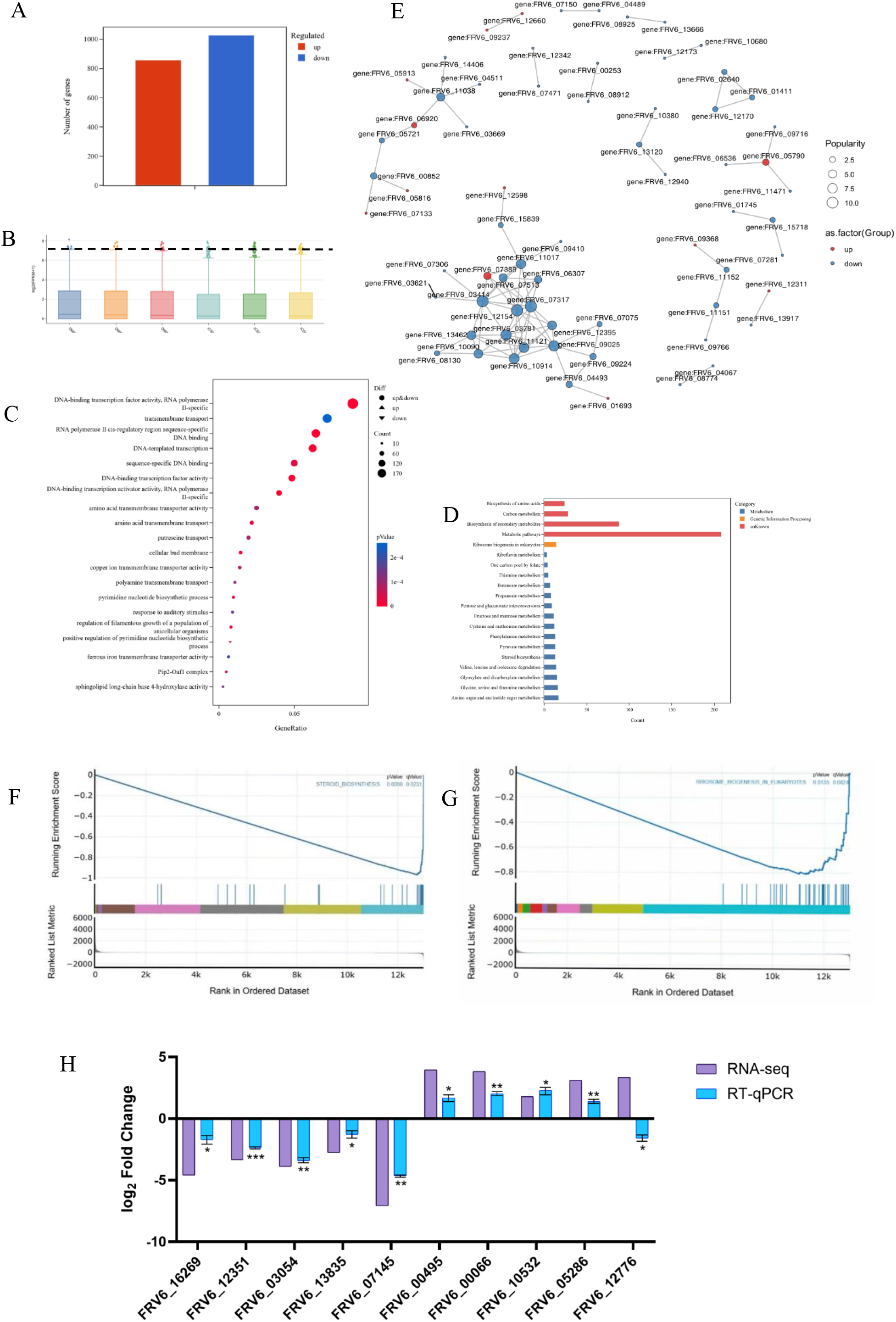
Transcriptomic profiling of *F. oxysporum* in response to D-limonene treatment. (A) Histogram of differentially expressed genes (DEGs). (B) Box plots of gene expression levels across samples. Each box represents five summary statistics (maximum, upper quartile, median, lower quartile, and minimum). The dashed line facilitates visual comparison between the treatment and control groups. (C) Bubble plot of GO enrichment. (D) Bar chart of KEGG pathway enrichment. (E) Protein–protein interaction network (PPIN) of DEGs. (F) Gene set enrichment analysis (GSEA) curve for steroid biosynthesis-related genes. (G) GSEA plot for ribosome biosyn-thesis-related genes. (H) Purple bars represent log₂FoldChange values obtained from RNA-seq analysis. Blue bars represent −ΔΔCt values from RT-qPCR (mean ±SEM, with 3 biological replicates). The asterisk (*) indicates a significant difference according to a t-test (\**P <* 0.05; \*\**P <* 0.01; \*\*\**P <* 0.001). In all figure panels, “limonene” refers specifically to D-limonene.

#### 3.9.2 Ergosterol Biosynthesis-Related Genes

To dissect the molecular basis underlying D-limonene-induced membrane damage, we focused on ergosterol biosynthesis genes. Gene set enrichment analysis (GSEA) of steroid syn-thesis-related genes revealed a significant overall downregulation trend (Figure 4F). KEGG analysis indicated that 18 of 28 steroid synthesis-related genes were downregulated, of which 11 were specifically involved in ergosterol synthesis (Table S1). Representative downregulated genes included *FRV6_13835* (encoding a cytochrome P450 protein), and *FRV6_01176* (a member of the *ERG4/ERG24* family) and *FRV6_08680* (a sterol methyltransferase-related gene), with fold decreases of approximately 6.8, 6.3, and 5.6, respectively. These results suggest that D-limonene may disrupt membrane integrity in *F. oxysporum* cells, at least in part, by downregulating ergosterol synthesis genes, potentially compromising membrane fluidity and function.

#### 3.9.3 Ribosome Biogenesis-Related Genes

In addition, ribosome biogenesis was also substantially suppressed. GSEA revealed a consistent global downregulation trend across all 42 genes annotated to this pathway (Figure 4G). Consistently, KEGG and PPIN analysis revealed that 59 of 63 ribosome-related genes were downregulated, including 26 genes involved in large ribosomal subunit (LSU) biogenesis (Table S2). Representative genes, including *FRV6_11098* (involved in 60S ribosomal subunit maturation), *FRV6_07306* (Urb2/Npa2 family protein), *FRV6_09184* (DEAD/DEAH-box helicase), and *FRV6_07317* (ribosomal protein L10), were downregulated by approximately 2.3–2.7-fold. Since proper LSU assembly is required for efficient ribosome formation and translation, suppression of these genes is likely to impair protein synthesis capacity, thereby contributing to the observed growth inhibition of *F. oxysporum*.

#### 3.9.4 Cell-Wall Synthesis and Remodeling-Related Genes

KEGG analysis showed significant enrichment of amino sugar and nucleotide sugar metabolism, indicating that the metabolism of precursors required for fungal cell-wall polysaccharide biosynthesis may be affected by D-limonene treatment (*P* < 0.05, Figure 4D). Based on this observation, cell-wall-related genes were further examined. A total of 31 high-confidence differentially expressed genes associated with chitin metabolism, β-glucan remodeling, mannan/mannoprotein metabolism, and cell-wall structure or assembly were identified, including 29 upregulated and two downregulated genes. These genes comprised nine chitin-related genes, seven β-glucan-remodeling genes, nine mannan- or mannoprotein-related genes, and six genes involved in cell-wall structure or assembly. Notably, all six identified chitin synthase genes were upregulated. Representative genes included *FRV6_13775* (chitin synthase 5), *FRV6_14590* (chitinase 1), *FRV6_09199* (endo-1,3(4)-β-glucanase), and *FRV6_10453* (cell-wall protein PhiA), with log₂FC values of 2.75, 3.96, 2.21, and 2.42, respectively (Table S3). These findings indicate that D-limonene induced extensive transcriptional remodeling of fungal cell-wall-related processes.

## 4. Discussion

Plant-derived and other naturally occurring bioactive substances have received increasing attention as alternatives or complements to conventional fungicides for the management of *Fusarium* diseases. These substances encompass several chemical classes, including terpenoids such as carvacrol, citral, L-menthol, and thymol;^39–41^ phenylpropanoid compounds represented by cinnamaldehyde;^40,42^ polysaccharide-derived compounds such as chitosan and chito-oligosaccharides;^34,43^ and other naturally occurring regulators, including melatonin.^44^ However, compared with other chemical classes, olefin-rich hydrocarbon monoterpenes have been investigated less extensively for their antifungal effects and mechanisms against *Fusarium* species. In the present study, D-limonene inhibited the mycelial growth of *F. oxysporum* in a concentration-dependent manner, with an IC₅₀ of 8.32 μL/mL. Although hydrocarbon monoterpenes such as limonene are generally considered to exhibit weaker antifungal activity than oxygenated or phenolic terpenes, this value demonstrates a moderate but definite concentration-dependent inhibitory effect against *F. oxysporum*. D-Limonene also induced pronounced changes in colony and hyphal morphology. More importantly, it markedly reduced spore germination and viability, suggesting that its antifungal activity extends beyond the suppression of established mycelial growth to interference with spore establishment and fungal propagation. The reduction in lesion development on potato tissues and the altered hyphal ultrastructure further indicate that D-limonene affects multiple stages of the fungal life cycle and disease development rather than acting solely as a mycelial growth inhibitor.

To elucidate the molecular basis underlying these phenotypic effects, transcriptomic analysis was performed. The resulting data suggest that the antifungal activity of D-limonene involves multiple cellular processes. The coordinated downregulation of genes associated with ergosterol biosynthesis provides a plausible molecular explanation for the loss of membrane integrity indicated by PI/FDA staining. Given the essential role of ergosterol in maintaining fungal membrane fluidity and function,^46^ suppression of its biosynthetic pathway may contribute to membrane dysfunction in D-limonene-treated *F. oxysporum*. This interpretation is also supported by previous reports describing membrane-disruptive effects of limonene and related monoterpenoids.^26,29,30,32,47,48^ In parallel, GSEA, KEGG enrichment, and PPIN analyses consistently indicated suppression of ribosome biogenesis, particularly the assembly of the large ribosomal subunit, suggesting that D-limonene reduces translational capacity. Comparable inhibition of ribosome-related processes has been reported in *F. graminearum* exposed to cinnamaldehyde or tea-derived bioactive mixtures,^42,49^ supporting ribosome biogenesis as a relevant antifungal vulnerability.^50^ Moreover, the altered FB 28 staining pattern, together with the predominant upregulation of genes involved in chitin synthesis and turnover, β-glucan remodeling, and cell-wall structure, is more consistent with a compensatory cell-wall stress response than with direct inhibition of cell-wall biosynthesis. Similar remodeling responses have been observed following limonene treatment in *C. albicans* and *C. tropicalis* and after chitosan or chito-oligosaccharide treatment in *F. oxysporum*.^26,29,34,43^ Collectively, these results support a multifaceted mode of action involving membrane dysfunction, suppression of ribosome biogenesis, and extensive remodeling of cell-wall-associated processes. Such multi-target activity may be advantageous for resistance management because it reduces reliance on a single cellular target and may complement strategies intended to mitigate the limitations of prolonged conventional fungicide use.^9,10^

The application potential of D-limonene was further supported by its interactions with conventional fungicides and environmental stresses. Previous studies have demonstrated that plant-derived volatile compounds or essential oils can enhance the efficacy of commercial fungicides, as reported for thymol combined with tebuconazole or difenoconazole,^47,48^ *Tagetes filifolia* essential oil combined with carbendazim, difenoconazole, or trifloxystrobin–cyproconazole,^51^ and plant essential oils combined with iprodione.^52^ These findings provide a basis for integrating natural products with synthetic fungicides. In the present study, D-limonene showed an additive interaction with mancozeb and a mildly synergistic interaction with hymexazol, resulting in greater growth inhibition than that achieved by either fungicide alone. This result suggests that D-limonene may serve as a complementary component of integrated PDR management and could potentially contribute to the optimization of fungicide inputs; however, any reduction in fungicide dose and maintenance of disease-control efficacy must be validated in tuber and storage trials. The stress-response assays also showed that D-limonene increased the sensitivity of *F. oxysporum* to temperature, salinity, and oxidative stresses, whereas its interaction with UV irradiation was limited. Although high temperature produced the strongest combined inhibition *in vitro*, low temperature is more directly relevant to potato storage, during which PDR commonly develops ^53^ and edible tubers are generally maintained at 2–4 °C.^54^ The enhanced inhibitory activity observed under low-temperature conditions therefore supports further evaluation of D-limonene as a storage-stage treatment. Nevertheless, formulation stability, tuber phytotoxicity, residue behavior, and efficacy under commercial storage conditions require additional investigation.

## 5. Conclusions

This study demonstrates that D-limonene exerts broad antifungal activity against *F. oxysporum*, a causal agent of potato dry rot. D-Limonene inhibited mycelial growth in a con-centration-dependent manner, with an IC₅₀ value of 8.32 μL/mL, and markedly reduced pathogenicity, spore germination, and spore viability while inducing pronounced alterations in colony and hyphal morphology. The redistribution and enhancement of fluorescent brightener 28 staining, together with the transcriptional activation of genes associated with chitin metabolism, β-glucan remodeling, mannan/mannoprotein metabolism, and cell-wall organization, indicate that D-limonene perturbs cell-wall homeostasis and elicits a compensatory remodeling response. Transcriptomic analyses further revealed coordinated suppression of ergosterol biosynthesis and ribosome biogenesis. These changes provide a molecular basis for the observed loss of membrane integrity and growth inhibition and suggest that limonene acts through a multifaceted mechanism involving membrane dysfunction, impaired translational capacity, and extensive cell-wall remodeling rather than through a single cellular target. D-Limonene also increased the sensitivity of *F. oxysporum* to temperature, salinity, and oxidative stresses and exhibited additive and mildly synergistic interactions with mancozeb and hymexazol, respectively. The enhanced inhibitory activity under low-temperature conditions is particularly relevant to postharvest potato storage. Collectively, these findings support D-limonene as a promising bio-based component of integrated potato dry rot management; however, its formulation stability, tuber safety, residue behavior, and efficacy under commercial storage conditions require further evaluation.

## Data Availability Statement

The datasets generated during and/or analysed during the current study are available from the corresponding author on reasonable request.

## Acknowledgments

We sincerely thank Professor Zhengguo Li at the School of Life Sciences, Chongqing University, China, for guidance on the methods and implementation of the experiments and for providing research funding. We would like to acknowledge that English language editing was partially assisted by ChatGPT and DeepSeek; all scientific content, data analysis and manuscript writing were independently completed by the authors.

## Supporting Information

Supporting information is provided in a separate file and includes Tables S1–S4 and Figure S1.

## References

1 Tiwari RK, Kumar R, Sharma S, Sagar V, Aggarwal R, et al., Potato dry rot disease: current status, pathogenomics and management. 3 Biotech 10:1–18 (2020).

2 Weiss F, Survey for potato wilt in Pennsylvania and Southern New York. Am J Potato Res 1:243–244 (1924).

3 Christian CL, Rosnow J, Woodhall JW, et al., Pathogenicity of *Fusarium* species associated with potato dry rot in the Pacific Northwest of the United States. Plant Dis 109:1091–1101 (2025).

4 Muratali D, Dervis S, Özer G, et al., Molecular and pathogenic characterization of *Fusarium* species associated with dry rot in stored potatoes in Kyrgyzstan. Potato Res 68:3271–3293 (2025).

5 Bojanowski A, Avis TJ, Pelletier S and Tweddell RJ, Management of potato dry rot. Postharvest Biol Technol 84:99–109 (2013).

6 Li Y, Xia X, Zhao Q and Dong P, The biocontrol of potato dry rot by microorganisms and bioactive substances: A review. Physiol Mol Plant Pathol 122:101919 (2022).

7 Sun Q, Li C and Yi X, The antibacterial effect of several biocontrol bacteria on *Fusarium oxysporum* and the rapid detection of their inclusive growth. Chin J Trop Agric 39:36–41 (2019).

8 Wang Y, Zhang G, Jiao J and Gao Z, The inhibition and disease control effect of 7 kinds of fungicides to potato dry rot fungus by *Fusarium solani*. Inner Mong Agric Sci Technol 43:83–85 (2015).

9 Platt H, Resistance to thiabendazole in *Fusarium* species and *Helminthosporium solani* in potato tubers treated commercially in eastern Canada. Phytoprotection 78:1–10 (1997).

10 Gurajala S and Challapilla M, A quiet but growing threat: Global perspectives on antifungal resistance and future therapeutic approaches. Indian J Pharmacol 58:35–45 (2026).

11 Lin H, Liu Y, Zhang X, et al., D-Limonene: Promising and sustainable natural bioactive compound. Appl Sci 14:4605 (2024).

12 Misra G, Pavlostathis SG, Perdue EM and Araujo R, Aerobic biodegradation of selected monoterpenes. Appl Microbiol Biotechnol 45:831–838 (1996).

13 Ravichandran C, Badgujar PC, Gunde VP, et al., Review of toxicological assessment of D-limonene, a food and cosmetics additive. Food Chem Toxicol 120:668–680 (2018).

14 Ibrahim MA, Kainulainen P, Aflatuni A, et al., Insecticidal, repellent, antimicrobial activity and phytotoxicity of essential oils: with special reference to limonene and its suitability for control of insect pests. Agric Food Sci Finl 10:243–259 (2001).

15 Hollingsworth RG, Limonene, a citrus extract, for control of mealybugs and scale insects. J Econ Entomol 98:772–779 (2005).

16 Zhang RF, Jin Y and Shi GL, Biological activity of D-limonene, the main component of orange peel. J Zhejiang Agric Sci 1:73–74 (2013).

17 Yang S, Study on fumigation activities of plant essential oil to stored grain pest. Thesis, Huazhong Agricultural University, Wuhan (2008).

18 Hebeish A, Fouda MMG, Hamdy IA, et al., Preparation of durable insect repellent cotton fabric: limonene as insecticide. Carbohydr Polym 74:268–273 (2008).

19 Zhang XX, Study on the activity of D-limonene in the control of *Panonychus citri* McGregor. Thesis, South China Agricultural University, Guangzhou (2018).

20 Badawy MEI, Abdelgaleil SAM, Mahmoud NF, et al., Preparation and characterization of essential oil and monoterpene nanoemulsions and acaricidal activity against two-spotted spider mite (Tetranychus urticae Koch). Int J Acarol 44:330–340 (2018).

21 Ellis MD and Baxendale FP, Toxicity of seven monoterpenoids to tracheal mites (Acari: Tarsonemidae) and their honey bee (Hymenoptera: Apidae) hosts when applied as fumigants. J Econ Entomol 90:1087–1091 (1997).

22 Abdelgaleil SAM, Mohamed MIE, Badawy MEI, et al., Fumigant and contact toxicities of monoterpenes to Sitophilus oryzae L, and *Tribolium castaneum* Herbst and their inhibitory effects on acetylcholinesterase activity. J Chem Ecol 35:518–525 (2009).

23 Isman MB, Miresmailli S and Machial C, Commercial opportunities for pesticides based on plant essential oils in agriculture, industry and consumer products. Phytochem Rev 10:197–204 (2011).

24 Gettys LA, Thayer KL and Sigmon JW, Phytotoxic effects of acetic acid and D-limonene on four aquatic plants. HortTechnology 32:110–118 (2022).

25 Patrício FFS, Simon CA, Silva MB, et al., D-limonene in the postharvest of ’THB’ papaya. Pesqui Agropecu Bras 59:e03426 (2024).

26 Thakre A, Zore G, Kodgire S, et al., Limonene inhibits Candida albicans growth by inducing apoptosis. Med Mycol 56:565–578 (2018).

27 Cai R, Hu M, Zhang Y, et al., Antifungal activity and mechanism of citral, limonene and eugenol against *Zygosaccharomyces rouxii*. LWT 106:50–56 (2019).

28 Leite-Andrade MC, de Araújo Neto LN, Buonafina-Paz MDS, et al., Antifungal effect and inhibition of the virulence mechanism of D-limonene against *Candida parapsilosis*. Molecules 27:8884 (2022).

29 Xiong HB, Zhou XH, Xiang WL, et al., Integrated transcriptome reveals that D-limonene inhibits *Candida tropicalis* by disrupting metabolism. LWT 176:114535 (2023).

30 Zhou SW, Zhu Y, Qin XJ, et al., D-limonene inhibits the growth of *Fusarium proliferatum* by decreasing H3K9ac and H3K27ac modifications. BMC Genomics 27:55 (2026).

31 Dambolena JS, Lopez AG, Canepa MC, et al., Inhibitory effect of cyclic terpenes (limonene, menthol, menthone and thymol) on *Fusarium verticillioides* MRC 826 growth and fumonisin B1 biosynthesis. Toxicon 51:37–44 (2008).

32 Jian YQ, Chen X, Ma HQ, et al., Limonene formulation exhibited potential application in the control of mycelial growth and deoxynivalenol production in *Fusarium graminearum*. Front Microbiol 14:1161244 (2023).

33 Marei GIK, Rasoul MAA and Abdelgaleil SAM, Comparative antifungal activities and biochemical effects of monoterpenes on plant pathogenic fungi. Pestic Biochem Physiol 103:56–61 (2012).

34 Ren J, Tong J, Li P, Huang X, Dong P and Ren M, Chitosan is an effective inhibitor against potato dry rot caused by *Fusarium oxysporum*. Physiol Mol Plant Pathol 113:101601 (2021).

35 Wang L, Xu T, Li L, et al., Biological characteristics and indoor fungicide screening of potato dry rot pathogen. J Xinjiang Agric Univ 39:222–226 (2016).

36 Ning N, Liu Q, Xian W, et al., Indoor toxicity determination of five fungicides against potato dry rot fungi. J Qinghai Univ 39:31–37 (2021).

37 Zhao D, Wei W, Zhang D, et al., Study on indoor fungicide screening and disease prevention of potato dry rot. Hubei Agric Sci 56:3268–3269, 3279 (2017).

38 Jin ZJ, About the evaluation of drug combination. Acta Pharmacol Sin 25:146–147 (2004).

39 Lu X, Hu K, Ou M, et al., Synergistic antifungal activity and mechanism of carvacrol/cit-ral combination against *Fusarium oxysporum* in Dendrobium officinale. Pestic Biochem Physiol 215:106671 (2025).

40 Kmoch M, Loubová V, Švecová R and Jílková B, Effect of essential oil components on the growth inhibition of *Fusarium solani* var. coeruleum during potato storage. Agronomy 15:1126 (2025).

41 Liu Y, Liu S, Lou X, et al., Antifungal activity and mechanism of thymol against *Fusarium oxysporum*, a pathogen of potato dry rot, and its potential application. Postharvest Biol Technol 192:112025 (2022).

42 Zhang C, Liu H, Wang X, et al., Inhibitory effects and mechanisms of cinnamaldehyde against *Fusarium oxysporum*, a serious pathogen in potatoes. Pest Manag Sci 80:3540–3552 (2024).

43 Huang A, Ren J, Zhang J, et al., A multi-pronged approach to understanding the antifungal mechanism of chito-oligosaccharide against *Fusarium oxysporum*, a major pathogen causing potato dry rot. Postharvest Biol Technol 232:114005 (2025).

44 Mbatha LA and Mbili NC, Effects of melatonin and Bacillus amyloliquefaciens MPA 1034 on the postharvest quality of potato tubers. Horticulturae 11:1119 (2025).

45 Song L, Wang S, Zou H, Yi X, Jia S, Li R and Song J, Regulation of ergosterol biosynthesis in pathogenic fungi: Opportunities for therapeutic development. Microorganisms 13:862 (2025).

46 Jafri H and Ahmad I, Combinational effect of essential oil compounds and antimicrobial drugs on *Candida albicans* and *Staphylococcus aureus* mixed biofilms. J Essent Oil Bear Plants 23:697–709 (2020).

47 Shcherbakova L, Mikityuk O, Arslanova L, et al., Studying the ability of thymol to improve fungicidal effects of tebuconazole and difenoconazole against some plant pathogenic fungi in seed or foliar treatments. Front Microbiol 12:629429 (2021).

48 Li Y, Wang X, Li H, et al., Mechanism of tea infusion as a novel control agent against *Fusarium graminearum*. Phytopathol Res 8:33 (2026).

49 Sun N, Li D, Zhang Y, Killeen K, Groutas W and Calderone R, Repurposing an inhibitor of ribosomal biogenesis with broad anti-fungal activity. Sci Rep 7:17014 (2017).

50 Gadban LC, Camiletti BX, Bigatton ED, Distéfano SG and Lucini EI, Combinations of Tagetes filifolia Lag, essential oil with chemical fungicides to control *Colletotrichum trunca-tum* and their effects on the biocontrol agent *Trichoderma harzianum*. J Plant Prot Res 60:41–50 (2020).

51 Camiletti BX, Asensio CM, Gadban LC, Pecci M de la PG, Conles MY and Lucini EI, Essential oils and their combinations with iprodione fungicide as potential antifungal agents against white rot (*Sclerotium cepivorum* Berk) in garlic (*Allium sativum* L.) crops. Ind Crops Prod 85:117–124 (2016).

52 Pooja, Chauhan P, Kumar A, Rithesh L and Kumar A, Recent insights into potato dry rot an emerging disease: Focusing on pathogen diversity, host-pathogen interactions, and management strategies. Microb Pathog 207:107866 (2025).

53 Rastovski A and van Es A, Storage of Potatoes: Post-Harvest Behaviour, Store Design, Storage Practice, Handling, Pudoc, Wageningen, The Netherlands (1987).

